# Restrictome-EVOLVE: population-resolved haplotype architecture of human antiviral restriction-factor loci

**DOI:** 10.64898/2026.09.01.748667

**Authors:** Reuben S. Maghembe, Samweli Y. Bahati, Abdalah Makaranga

## Abstract

Human antiviral restriction factors act across multiple stages of viral replication, but whether their population-resolved haplotype architecture differs systematically from comparable genomic regions is unclear. We tested this using phased public human genomic data from 660 individuals in seven African and African-diaspora populations, representing 30 canonical restriction-factor units and 436 target windows. Each canonical unit was compared with 80 exact matched genomic controls, yielding 2,400 frozen controls and 4,429,760 target-control endpoint comparisons across 19 retained haplotype endpoints. All 30 canonical units showed lower differentiation effects and lower robust population-private haplotype effects than their matched controls. Within-population diversity effects were higher in 19 of 30 units, whereas dominant-haplotype concentration effects were lower in 22 of 30. Nineteen units occupied a deconcentrated/high-diversity state, eight a concentrated/low-diversity state, and three a lower-diversity/lower-concentration state. Of 127 global endpoint/context summaries, 88 reached a global false-discovery-rate q value below 0.05; 76 were lower in restriction-factor targets and 12 were higher. Directional sign-test inference detected widespread repeated displacement relative to matched controls, whereas no matched-cell empirical-rank test reached global false-discovery-rate significance. These results show that human antiviral restriction-factor loci occupy a reproducible matched-control haplotype architecture characterized by attenuated population partitioning and reduced robust private structure, together with substantial locus-specific variation in within-population diversity and haplotype concentration. The comparative framework separates population-genomic structure from claims of functional or adaptive causality.

## Introduction

Human antiviral restriction factors are host-encoded molecules that can interfere with viral entry, genome release, replication, intracellular trafficking, assembly, egress, cell-to-cell spread, or persistence. They include membrane-associated proteins, nucleic-acid sensors, deaminase-related enzymes, interferon-stimulated effectors, RNA-binding proteins, and intracellular trafficking regulators. Rather than forming a single linear pathway, these proteins constitute a distributed and context-dependent defense system whose activity depends on viral family, tissue and cell state, interferon exposure, viral antagonists, and host genetic background (Chemudupati et al. 2019; Duggal and Emerman 2012; Sadler and Williams 2008; Schneider et al. 2014). Restriction-factor genes are therefore natural targets for evolutionary and population-genomic analysis. Host-defense loci are shaped jointly by mutation, recombination, demographic history, drift, admixture, functional constraint, and—in some contexts—pathogen-mediated selection. Molecular conflict between host restriction factors and viral antagonists can further generate unusual evolutionary trajectories, while population immune-genomic studies have shown that regulatory and coding variation can contribute to differences in immune-response phenotypes (Barreiro and Quintana-Murci 2010; Daugherty and Malik 2012; Enard et al. 2016; Karlsson et al. 2014; Nedelec et al. 2016; Quach et al. 2016; Quintana-Murci and Clark 2013; Sironi et al. 2015). Importantly, however, population differentiation alone does not identify the evolutionary process responsible for a signal and does not establish functional or clinical consequences. Much population-genomic interpretation is performed at the level of individual variants or allele-frequency differences. For restriction-factor loci, a complementary level of information is carried by phased haplotypes. Haplotype distributions capture combinations of linked alleles, the concentration or dispersion of common configurations, population-private structures, and differences in the overall frequency distribution among populations. These properties can reveal whether a locus is unusually partitioned among populations, unusually concentrated within populations, or unusually rich in distinct haplotypes. A central interpretive challenge is that the same patterns can also arise from local genomic properties and demographic history. Restriction-factor loci therefore require comparison with genomic regions that are matched as closely as possible for relevant background structure.

Here we tested this comparative question by analysing phased haplotype architecture across seven African and African-diaspora populations and organized the antiviral gene framework into 30 canonical physical units. Each unit was compared with 80 exact matched genomic controls across a retained panel of differentiation, population-private, diversity, concentration, grouped, cross-population, and omnibus endpoints. We asked whether restriction-factor loci occupy distinctive positions relative to their matched genomic background, while explicitly separating repeated directional displacement from empirical extremeness within the control distribution. The framework shifts the emphasis from isolated candidate variants to the population-resolved architecture of antiviral restriction-factor haplotypes.

## Results

### Restriction-factor loci show coordinated attenuation of population partitioning relative to matched genomic controls

We compared phased haplotype architecture across 30 canonical restriction-factor units with structurally and population-aware matched genomic controls. A consistent matched-control pattern emerged across the restriction-factor panel: all 30/30 canonical units had negative median standardized differentiation effects relative to their matched controls (Fig 1A). The same directionality extended to robust population-private haplotype structure, for which all 30/30 units also showed negative effects. Thus, the dominant locus-level signature was not increased partitioning among the sampled populations, but broad attenuation of both between-population differentiation and robust private-haplotype structure relative to comparable genomic regions.

**Fig 1.**
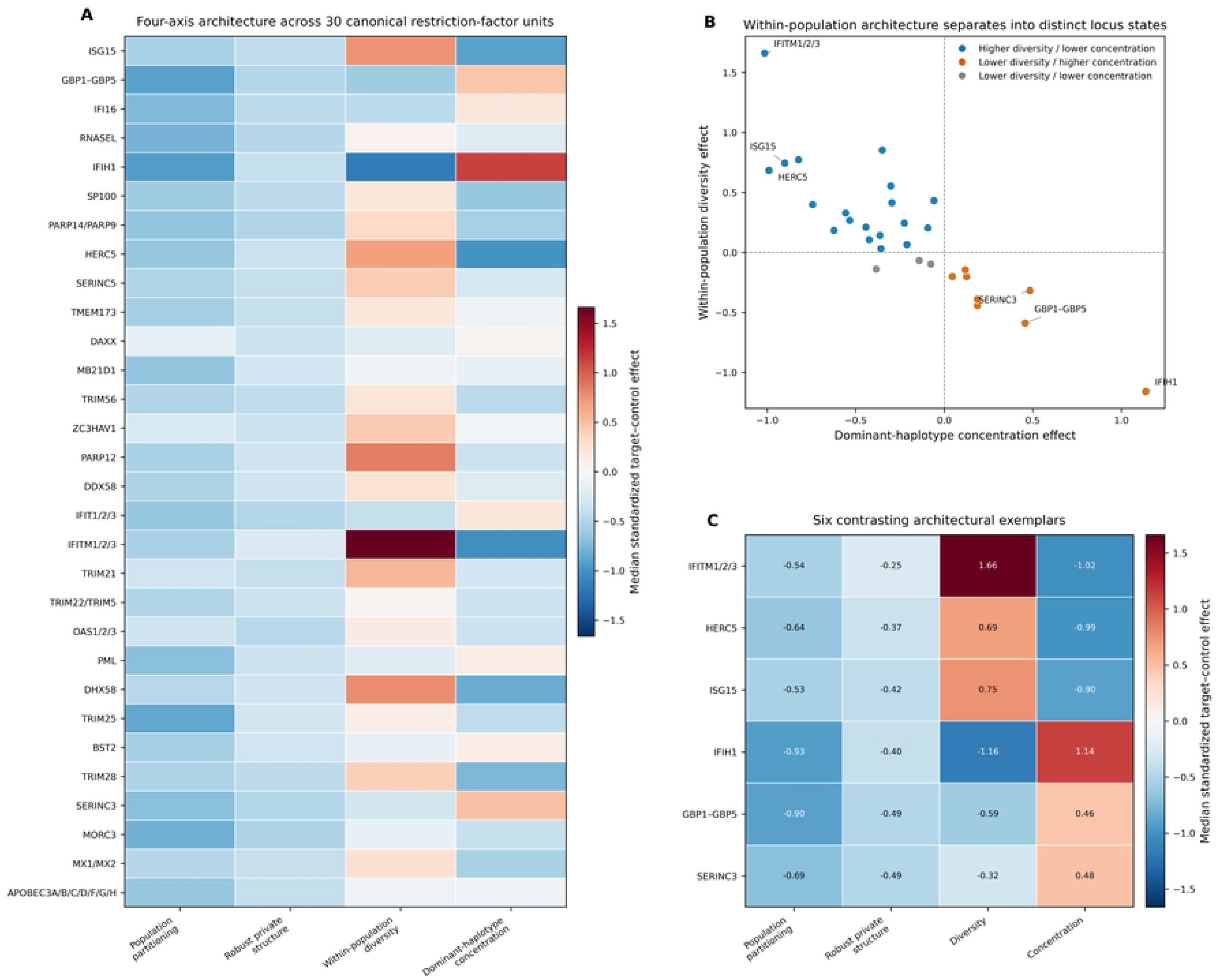
Restriction-factor loci share reduced population partitioning but diverge in within-population architecture. (A) Four-axis architecture across the 30 canonical restriction-factor units. Cells show the median standardized target-minus-control effect for population differentiation, robust private-haplotype structure, within-population haplotype diversity, and dominant-haplotype concentration. Negative values indicate lower values in the restriction-factor target unit relative to its matched genomic controls; positive values indicate higher values in the target. Gene labels denote the canonical physical units used throughout the analysis. (B) Relationship between the canonical-unit within-population diversity effect and dominant-haplotype concentration effect. The observed units occupy three sign-defined architectural states: higher-diversity/lower-concentration, lower-diversity/higher-concentration, and lower-diversity/lower-concentration. Six extreme architectural exemplars are labelled. (C) Four-axis profiles of the six contrasting exemplar units selected from panel B: the three strongest deconcentrated/high-diversity and the three strongest concentrated/low-diversity architectures. Numeric values are median standardized target-minus-control effects. All panels were generated from the frozen matched-control comparative inference layer. Color scales are centered at zero; blue denotes lower and red denotes higher values in restriction-factor targets relative to matched genomic controls.

The four-axis canonical-unit representation nevertheless showed that this shared attenuation of population partitioning did not imply a uniform within-population haplotype configuration (Fig 1A). Within-population diversity effects were positive in 19/30 units, whereas dominant-haplotype concentration effects were negative in 22/30. The combination of these axes therefore separated a common between-population signal from substantial locus-specific variation in the organization of haplotypes within populations.

### Within-population haplotype architecture separates restriction-factor loci into contrasting states

Joint consideration of diversity and top-haplotype concentration identified three recurrent architectural states (Fig 1B). The predominant configuration was deconcentrated/high-diversity (19/30 units), followed by concentrated/low-diversity (8/30) and lower-diversity/lower-concentration (3/30). This distribution indicates that reduced population partitioning can coexist with markedly different local haplotype states.

The six exemplar loci highlighted in Fig 1C capture the extremes of this continuum. The deconcentrated/high-diversity exemplars were *IFITM1*/*IFITM2*/*IFITM3*, *HERC5*, *ISG15*, whereas *IFIH1*, *GBP1*/*GBP2*/*GBP3*/*GBP4*/*GBP5*, *SERINC3* represented the strongest concentrated/low-diversity examples in the frozen canonical-unit matrix. These loci retained the common negative differentiation and robust-private axes while diverging most strongly in within-population diversity and dominant-haplotype concentration.

### Attenuated pairwise differentiation is widespread across African and diaspora population contrasts

For haplotype FST, 20/21 population-pair contrasts reached global FDR q < 0.05; 18 of these were lower in restriction-factor targets and 2 were higher. The strongest lower-in-target contrast was GWD–LWK (median standardized effect -1.22, global q = 1.18 × 10⁻⁸).

For Jensen–Shannon divergence, 15/21 population-pair contrasts reached global FDR q < 0.05, including 14 lower-in-target contrasts and one higher-in-target contrast. The strongest lower-in-target contrast was GWD–LWK (median standardized effect -0.82, global q = 1.83 × 10⁻⁷).

For maximum haplotype-frequency difference, 16/21 population-pair contrasts reached global FDR q < 0.05, and all 16 significant contrasts were lower in restriction-factor targets. The strongest lower-in-target contrast was ACB–GWD (median standardized effect -1.02, global q = 1.18 × 10⁻⁸).

Across the three pairwise metrics, the predominance of negative effects was visible across continental African and African-diaspora comparisons rather than being confined to a single population pair (Fig 2A-2C). The three metrics quantify related but distinct properties of haplotype-distribution separation, yet converged on the same overall pattern of attenuated population differentiation relative to the matched genomic background.

**Fig 2.**
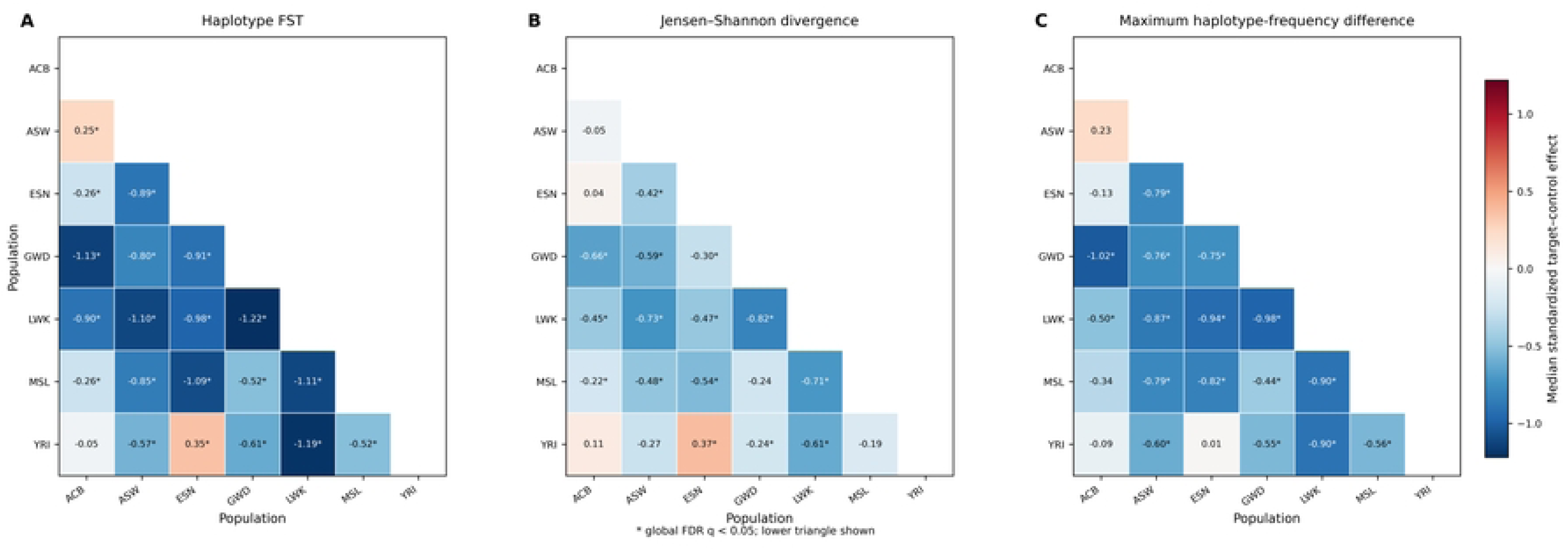
Restriction-factor loci show broadly attenuated haplotype differentiation across African and diaspora populations. Pairwise population-differentiation architecture across ACB, ASW, ESN, GWD, LWK, MSL, and YRI. (A) Haplotype FST. (B) Jensen–Shannon divergence between population haplotype-frequency distributions. (C) Maximum haplotype-frequency difference. Each displayed cell is the median standardized target-minus-control effect across the 30 canonical restriction-factor units for the corresponding population pair. The lower triangle is shown. Blue indicates reduced differentiation in restriction-factor targets relative to matched genomic controls and red indicates increased differentiation. Asterisks denote global FDR q < 0.05 from the across-unit directional sign-test framework. All three panels share the same standardized-effect color scale.

### Population-private structure is reduced while within-population diversity is selectively enriched

Population-level private-haplotype summaries further separated raw haplotype occurrence from robust private structure (Fig 3A). Robust private-haplotype counts and robust private-haplotype frequency mass were predominantly negative relative to matched controls across populations, consistent with the canonical-unit robust-private axis. Raw private-haplotype counts were more heterogeneous, demonstrating that an increased number of nominally private haplotypes need not translate into an increased burden of robust population-private structure.

**Fig 3.**
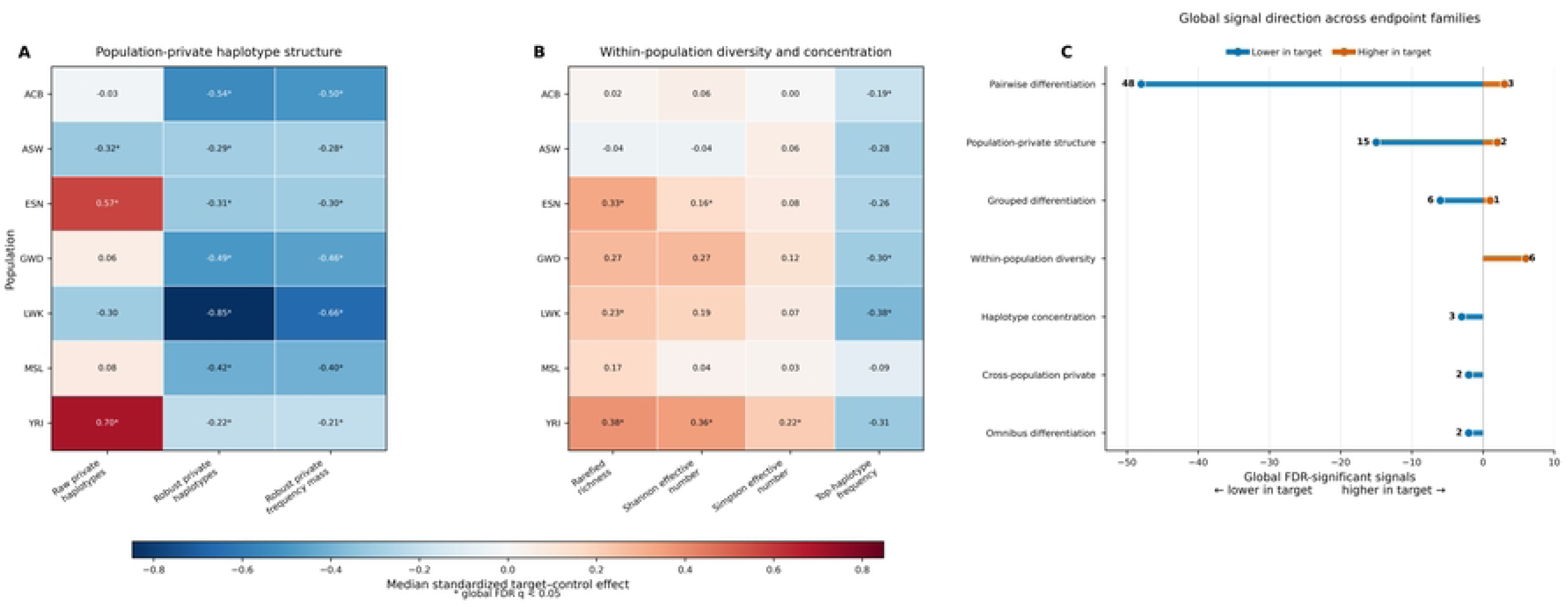
Private-structure attenuation coexists with selective within-population haplotype diversity enrichment. (A) Population-private haplotype architecture. Cells show median standardized target-minus-control effects for raw private-haplotype count, robust private-haplotype count, and robust private-haplotype frequency mass in each population. (B) Within-population haplotype diversity and concentration. Displayed metrics are expected rarefied haplotype richness, Shannon effective haplotype number, Simpson effective haplotype number, and top-haplotype frequency. Panels A and B share a zero-centered standardized-effect color scale. Asterisks denote global FDR q < 0.05 from the across-unit directional sign-test framework. (C) Direction and endpoint-family composition of the 88 globally significant endpoint/context signals. Blue lollipops denote the 76 lower-in-target signals and orange lollipops denote the 12 higher-in-target signals. Numbers adjacent to each endpoint family give the corresponding signal count. Only endpoint/context rows with global FDR q < 0.05 are included.

YRI provided the clearest higher-in-target raw-private example, with a median standardized effect of 0.70 and global q = 2.30 × 10⁻⁶, whereas the corresponding robust-private summaries remained negative (Fig 3A).

Within-population diversity metrics showed the complementary pattern (Fig 3B). Rarefied richness and effective haplotype-number measures were selectively elevated, whereas top-haplotype frequency was lower in several populations, producing a deconcentrated diversity profile. Representative globally significant higher-in-target signals included YRI rarefied richness (0.38, q = 1.20 × 10⁻⁴); YRI Shannon effective haplotype number (0.36, q = 6.07 × 10⁻⁴); ESN rarefied richness (0.33, q = 0.009).

### Global comparative inference is dominated by lower-in-target endpoint signals

Across the complete global endpoint/context synthesis, 88 of 127 rows reached global FDR q < 0.05. Of these significant signals, 76 were lower in restriction-factor targets and 12 were higher (Fig 3C). The significant set was therefore strongly asymmetric toward lower-in-target effects. The directional sign-test and matched-control empirical-rank tests captured distinct aspects of the comparative architecture. Directional displacement was widespread at the matched-cell level (44,809 global-FDR discoveries; 44,688 endpoint/context-FDR discoveries), whereas no matched-cell empirical-rank test reached global FDR q < 0.05. Thus, the principal signal reflects repeated directional displacement of restriction-factor targets relative to their matched controls rather than universal extremeness of targets within the corresponding matched-control distributions.

Pairwise population differentiation contributed the largest share of the significant signal architecture (51/88 signals), followed by population-private haplotype structure (17), grouped population differentiation (7), within-population haplotype diversity (6), haplotype concentration (3), cross-population private structure (2), and omnibus population differentiation (2).

Primary-tier endpoints accounted for 76 of the 88 significant signals, whereas 12 arose from secondary-tier endpoints. Taken together, the global synthesis identifies a restriction-factor haplotype architecture characterized by broad attenuation of population partitioning and robust private structure, superimposed on selective enrichment of within-population haplotype diversity at a subset of loci and populations.

## Discussion

Restrictome-EVOLVE identifies a reproducible comparative haplotype architecture across antiviral restriction-factor loci. All 30 canonical units showed negative differentiation effects relative to matched genomic controls, and all 30 also showed negative robust population-private effects. The concordance of these two axes is informative because they describe related but non-identical aspects of population structure. Together they indicate that, relative to genomic regions selected under the matching framework, the analysed restriction-factor loci tend to be less strongly partitioned among the sampled populations and contain less robust population-private haplotype structure.

This pattern should not be interpreted as evidence that restriction-factor loci are genetically homogeneous. Instead, it defines their position relative to the matched genomic background. Restriction factors are embedded in a biological landscape shaped by host-virus interaction, demographic history, recombination, constraint, and potentially multiple forms of selection (Barreiro and Quintana-Murci 2010; Daugherty and Malik 2012; Duggal and Emerman 2012; Enard et al. 2016; Karlsson et al. 2014; Sironi et al. 2015). The present design does not attempt to assign those processes individually. Its contribution is to establish a reproducible population-resolved comparative architecture against which more specific evolutionary hypotheses can subsequently be tested.

Pairwise population differentiation accounted for the largest component of the global significant signal set. Fifty-one of the 88 globally significant endpoint/context rows arose from pairwise differentiation, and the three principal pairwise metrics—haplotype FST, Jensen-Shannon divergence, and maximum haplotype-frequency difference—were dominated by lower-in-target effects. This convergence is important because the metrics summarize different properties of the haplotype-frequency distribution. Their agreement therefore argues that the observed attenuation is not restricted to a single mathematical representation of population separation.

The strongest negative contrasts were not confined to one population pair. Instead, lower-in-target effects extended across multiple comparisons involving continental African and African-diaspora populations. Selected higher-in-target exceptions remained, demonstrating that the architecture is not uniform. The appropriate biological interpretation is therefore one of broad attenuation superimposed on contrast-specific heterogeneity, rather than a claim that every restriction-factor locus or every population comparison follows the same evolutionary trajectory. The population-private analyses add an important distinction. Robust private-haplotype counts and robust private-haplotype frequency mass were predominantly reduced relative to matched controls, and the canonical-unit robust-private axis was negative in all 30 units. Raw private-haplotype counts were more heterogeneous and included higher-in-target signals. YRI, for example, showed a prominent positive raw-private signal while the corresponding robust-private measures remained negative.

This separation indicates that the mere observation of a haplotype only in one sampled population is not equivalent to evidence for a stable, population-private haplotype structure. Rare configurations are sensitive to sampling and frequency thresholds, whereas robust-private summaries place greater emphasis on private haplotypes with stronger frequency support. The present results therefore argue for explicit separation of raw private occurrence from robust private structure in population-genomic analyses of antiviral loci.

Reduced between-population partitioning coexisted with substantial within-population diversity. Nineteen of 30 canonical units showed positive diversity effects, and 22 of 30 showed reduced dominant-haplotype concentration. The predominant locus-level state was therefore deconcentrated/high-diversity rather than simple haplotype depletion. Population-level signals were consistent with this interpretation: selected richness and effective-haplotype-number endpoints were elevated, while top-haplotype frequency was reduced in several contexts.

This combination is biologically different from both a low-diversity selective-sweep-like pattern and a strongly population-partitioned architecture. It is compatible with multiple evolutionary and demographic histories, including recombination, long-term maintenance of alternative haplotypes, weakly structured variation, or heterogeneous locus-specific constraint. The present matched-control analysis cannot distinguish among these mechanisms, but it narrows the empirical pattern that any future evolutionary explanation must account for.

The four-axis canonical-unit analysis demonstrates that the global pattern does not erase locus-specific structure. Nineteen units occupied a deconcentrated/high-diversity state, eight a concentrated/low-diversity state, and three a lower-diversity/lower-concentration state. *IFITM1*/2/3, *HERC5*, and *ISG15* were among the strongest deconcentrated/high-diversity examples, whereas *IFIH1*, the *GBP1*-5 cluster, and *SERINC3* represented strong concentrated/low-diversity examples.

These exemplars are best interpreted as architectural representatives, not as demonstrations of gene-specific causal mechanisms. Restriction factors operate through diverse antiviral processes (Agol and Gmyl 2010; Chemudupati et al. 2019; Colomer-Lluch et al. 2018; Duggal and Emerman 2012; Everett 2018; Hatziioannou and Bieniasz 2011; Laguette et al. 2011; Lin et al. 2025; Sadler and Williams 2008; Schneider et al. 2014), and the same population-genomic pattern could arise through different combinations of molecular constraint, demographic history, recombination, regulatory variation, or past host-virus conflict. The value of the locus-level classification is therefore to identify distinct genomic states for downstream functional and evolutionary testing.

The seven-population design was deliberately population resolved rather than collapsed into broad continental superpopulations. This makes it possible to detect contrast-specific architecture while avoiding the assumption that a large umbrella ancestry category is biologically homogeneous. Population immune-genomic studies have previously shown that demographic history and genetic variation can influence immune-response phenotypes (Nedelec et al. 2016; Quach et al. 2016; Quintana-Murci and Clark 2013), but those observations do not imply deterministic immune characteristics for populations or individuals.

The present analysis should therefore be read at the level at which it was performed: as a comparison of haplotype distributions among defined public genomic population samples. The results do not establish differential antiviral protection, disease susceptibility, or clinical outcome among populations. Extending the work toward phenotype will require independent cohorts, explicit environmental and demographic modelling, and direct functional measurements.

A critical inferential feature of the study is the distinction between the directional sign-test and the empirical-rank analysis. The matched-cell sign-test layer produced 44,809 global-FDR discoveries and 44,688 endpoint/context-FDR discoveries, whereas no empirical-rank cell test survived global FDR correction. These findings are not contradictory.

The sign-test framework asks whether the target is repeatedly displaced in one direction across its 80 matched controls. A target can satisfy this condition while still occupying a non-extreme position within the full control distribution. Empirical rank asks the latter question directly. The combined result therefore supports widespread directional organization of restriction-factor haplotype architecture relative to matched genomic backgrounds, while providing no global-FDR evidence that individual target cells are generally extreme empirical outliers. Maintaining this distinction is essential for avoiding overinterpretation of the large sign-test discovery count. Restriction factors are often discussed through the framework of molecular conflict, because viral proteins can antagonize host restriction mechanisms and host proteins may evolve in response (Daugherty and Malik 2012; Duggal and Emerman 2012; Enard et al. 2016; Sironi et al. 2015). At the same time, population-genomic variation reflects demographic history, drift, recombination, admixture, structural variation, and functional constraint as well as selection (Barreiro and Quintana-Murci 2010; Karlsson et al. 2014; Quintana-Murci and Clark 2013). The present results are therefore compatible with several evolutionary histories and should not be treated as direct evidence of pathogen-driven positive selection, balancing selection, or local adaptation.

The matched-control design nevertheless provides a useful foundation for future evolutionary analysis because it identifies which aspects of restriction-factor haplotype architecture differ systematically from comparable genomic regions. Formal selection statistics, genealogical modelling, local-ancestry analysis, ancient-DNA or archaic-introgression analysis where appropriate, and population-specific recombination models can now be directed toward loci with clearly defined comparative architectures rather than selected solely from variant-level frequency differences.

Several design features support the robustness of the study. The analysis uses phased, population-resolved haplotypes; explicitly defined canonical physical units; exact frozen genomic controls; multiple complementary population endpoints; independent inference validation; and a checksum-locked manuscript-facing evidence layer. The 80-control design allows each target to be interpreted against a local matched distribution rather than a single comparator, while the separation of sign-test and empirical-rank inference prevents two distinct statistical questions from being conflated.

Important limitations remain. The analysis uses public population-genomic data and does not include direct viral infection phenotypes, clinical outcomes, cell-state-specific expression, chromatin accessibility, proteomic measurements, or experimental restriction assays. The seven populations do not represent the full diversity of African or global human populations. Complex structural variants, long repetitive regions, population-specific recombination histories, and local ancestry may not be fully captured by the present framework. Matched genomic controls reduce confounding by measured genomic characteristics but cannot guarantee exchangeability for every unmeasured evolutionary property.

Future work should therefore combine this architecture with formal selection analyses, demographic and genealogical modelling, regulatory genomics, allele-specific expression, long-read haplotypes, structural variation, and independent population cohorts. Functional studies can then test whether the contrasting architectural states identified here are associated with changes in restriction-factor expression, protein function, viral antagonism, or cellular antiviral activity. In this way, Restrictome-EVOLVE provides a population-genomic framework for generating mechanistically testable hypotheses while maintaining a clear boundary between comparative genomic structure and causal antiviral function.

## Materials and methods

### Study design and analytical framework

Restrictome-EVOLVE was conducted as a population-resolved, matched-genomic-control analysis of phased haplotype architecture across human antiviral restriction-factor loci. The final analytical panel comprised 660 individuals from seven 1000 Genomes populations: ACB (n=96), ASW (n=61), ESN (n=99), GWD (n=113), LWK (n=99), MSL (n=85), and YRI (n=107). Restriction-factor loci were represented by 30 canonical physical units and 436 refined target windows. Each canonical unit was paired with 80 matched genomic controls selected before endpoint testing, giving 2,400 final control regions. Population-resolved haplotype distributions were then quantified across differentiation, private-structure, diversity, concentration, grouped, cross-population, and omnibus contexts. Target values were interpreted relative to their matched-control distributions rather than against an unmatched genome-wide background.

The analytical design separated genomic matching, haplotype construction, endpoint computation, comparative inference, and manuscript-level synthesis. Control selection was therefore completed without reference to downstream haplotype endpoint values, and the inferential procedures were applied only after the target-control architecture had been fixed.

### Restriction-factor framework and canonical physical units

The antiviral restriction-factor framework was represented by 30 canonical physical units. Each unit corresponded either to a single restriction-factor locus or to a compact physically linked cluster whose constituent genes were analysed jointly under the final window architecture. This physical-unit representation provided a stable level for window construction, genomic-control matching, canonical-unit synthesis, and publication-level interpretation while avoiding double counting of overlapping or tightly linked loci.

The final canonical-unit architecture contained 436 refined target windows. Gene assignments and physical-unit definitions are reported in S1 Table. The canonical-unit framework, rather than individual candidate variants, was the primary organizational level of the present analysis.

### Population panel and phased genomic data

Genotypes were derived from the 1000 Genomes phase-three population resource (1000 Genomes Project Consortium 2015) and specifically from the 2019-03-12 integrated, phased, biallelic SNV and INDEL call set generated directly on the GRCh38 assembly (Lowy-Gallego et al. 2019). The study input was the matched-sample target extraction from this same phased GRCh38 release, and matched genomic controls were obtained from chromosome-level subsets of the identical source resource. The later 30x NYGC resource was not used as the source call set for the present analysis.

Population assignments were resolved before haplotype computation. The seven retained populations contributed 192 ACB, 122 ASW, 198 ESN, 226 GWD, 198 LWK, 170 MSL, and 214 YRI haploid chromosomes, respectively. Only complete phased genotype states within the selected analysis panels contributed to haplotype encoding. Each diploid individual contributed two phased haplotypes, which were encoded as ordered allele vectors within each analysis window and counted to obtain population-specific and pooled haplotype-frequency distributions.

### Haplotype windows and population LD portability

Window construction and tag selection were evaluated for portability across the seven populations before the full matched-control comparison. Of the 436 refined target windows, 431 supported the population-portable global tag-haplotype model. The remaining five windows were retained rather than excluded: three were represented with complex anchor-centred haplotype models and two with compact-tag haplotype models. Thus, all 436 refined windows entered downstream haplotype analysis, with model form adapted only where the global portable representation was not adequate.

Portable target windows used their validated global tag panels. The five complex windows used their prespecified alternative compact or anchor-centred representations. The corresponding target and control windows were evaluated under the fixed downstream panel contract so that comparative inference operated on aligned analytical entities rather than on post hoc lead-variant proxy expansions.

### Matched genomic control construction and exact control freeze

Candidate control regions were assembled outside the restriction-factor target intervals and evaluated using a multistage genomic matching design. The final matching objective combined population-aware genomic features and structural prematching information, with 80% of the optimization weight assigned to the population-aware component and 20% to structural prematching quality. The final feature contract comprised 61 matching variables. Selection also enforced the fixed spatial integrity constraints of the control design, including absence of target overlap, within-unit pairwise non-overlap, and the prespecified broad spatial-bin restriction.

Exact optimization yielded 80 controls for each of the 30 canonical units, for a total of 2,400 matched genomic controls. These assignments were fixed before haplotype endpoint evaluation. Each target refined window was subsequently linked to the appropriate control entities for comparative analysis, and every evaluable target endpoint/context cell was compared with exactly 80 matched-control values.

### Population-resolved haplotype endpoint computation

The retained analytical layer contained 19 haplotype endpoints spanning complementary properties of haplotype-frequency distributions. For a frequency vector p = (pᵢ), haplotype concentration was calculated as C = Σᵢ pᵢ² (Simpson 1949). Finite-sample haplotype diversity was calculated as D = [n/(n − 1)](1 − C) for n > 1 (Hurlbert 1971; Simpson 1949). Shannon entropy was H = −Σᵢ pᵢ ln(pᵢ) (Shannon 1948); the corresponding Shannon effective haplotype number was exp(H), and the Simpson effective haplotype number was 1/C, following the effective-diversity framework (Hill 1973).

Expected rarefied haplotype richness was calculated without replacement at a common depth of 122 haploid chromosomes, the smallest population chromosome count in the seven-population panel. For haplotype i with count nᵢ in a population of N chromosomes, expected richness at depth m was the sum across haplotypes of 1 − choose(N − nᵢ, m)/choose(N, m), implemented numerically in log-gamma form (Hurlbert 1971).

Population-distribution separation was described using several complementary measures. Total-variation distance was 0.5 times the sum of absolute frequency differences. Jensen–Shannon divergence used the symmetric base-2 formulation with the midpoint distribution m = (p + q)/2 (Lin 1991). Hellinger distance was calculated as the square root of one half of the summed squared differences between square-root frequencies. Pairwise and omnibus haplotype FST used the heterozygosity-ratio form max[0, (H◻ − H◻)/H◻], with H◻ weighted by the relevant population chromosome counts; this implementation follows the H◻/H◻ gene-diversity framework (Nei 1973) and is not the Weir–Cockerham theta estimator. Maximum haplotype-frequency difference supplied an additional direct measure of population separation.

Private-haplotype endpoints distinguished nominal private occurrence from robust population-private structure. A haplotype was considered robustly private only when it was private to one focal population, observed at least three times, and had within-population frequency ≥ 0.01. Raw private counts, robust private counts, and robust private frequency mass were therefore retained as distinct quantities. The endpoint quantities were instantiated across population-specific, pairwise, grouped, cross-population, and omnibus contexts to produce the 19 retained analytical endpoint definitions. Across targets and controls, the resulting endpoint layer contained 4,485,132 rows.

### Matched-control comparative inference

Comparative inference was performed within cells defined by canonical unit, refined target window, endpoint, analytical context, and target entity. A total of 55,372 evaluable target cells were reconstructed, each with exactly 80 matched controls, yielding 4,429,760 target-control comparisons.

The endpoint-specific transformation fixed in the analytical workflow was applied before target-control differencing. For transformed differences Δ, the standardized mean effect was mean(Δ)/SD(Δ) when SD(Δ) > 0. A robust complementary summary was median(Δ)/(1.4826 × MAD(Δ)) when MAD(Δ) > 0. Raw and transformed means and medians, standardized effects, robust effects, and directional concordance were retained for synthesis.

Primary cell-level directional inference used an exact two-sided sign test (Dixon and Mood 1946). Positive and negative transformed target-minus-control differences were counted, zero differences were treated as ties and omitted from the effective n, and the two-sided binomial tail probability was computed exactly. This test asks whether a target is repeatedly displaced in one direction across its matched controls.

A separate empirical-rank procedure evaluated target extremeness within the matched-control distribution. For the target transformed value and its 80 matched-control transformed values, lower and upper empirical tails were calculated with a +1 finite-reference correction, and the two-sided empirical probability was 2 times the smaller corrected tail probability, truncated at 1. This procedure was study-defined and source-locked. It answers a different question from the sign test: repeated directional displacement does not require that the target occupy an extreme rank within the control distribution.

### Multiplicity control and global synthesis

Multiple-testing correction used the Benjamini-Hochberg false-discovery-rate procedure (Benjamini and Hochberg 1995). At the matched-cell level, sign-test and empirical-rank probabilities were adjusted globally and within the prespecified analysis-tier/endpoint/context families. Canonical-unit synthesis combined refined-window effects while preserving equal canonical-unit contribution, and its inferential probabilities were adjusted globally and within analysis tier. Global endpoint/context synthesis similarly aggregated canonical-unit effects and applied the corresponding global and tier-specific correction structure.

The validated cell-level results comprised 44,809 sign-test discoveries at global FDR q < 0.05 and 44,688 at endpoint/context FDR q < 0.05. No empirical-rank cell reached global FDR q < 0.05. Canonical-unit synthesis produced 3,810 rows, including 1,216 global-FDR-significant rows, while the global endpoint/context layer contained 127 rows, of which 88 reached global FDR q < 0.05. The sign-test and empirical-rank results were therefore retained as complementary inferential layers rather than combined into a single significance interpretation.

### Validation, reproducibility, and publication-source generation

The global endpoint assembly and comparative-inference outputs were independently reconstructed and subjected to numerical, row-count, control-cardinality, checksum, and inferential validation before manuscript synthesis. Validation confirmed 55,372 inferential cells, exactly 80 controls per cell, 4,429,760 target-control comparisons, and the manuscript-level significance counts reported above. Analytical source files, implementation contexts, exact formulas, genomic-control contracts, callset provenance, and software/environment evidence were retained in the reproducibility package.

The final software record includes the versioned Python scientific-computing and genomic-tool environment used by the frozen analyses, including NumPy, pandas, bcftools, bedtools, PLINK/PLINK2, and tabix. Publication figures were generated in 600-DPI TIFF, vector PDF, and editable SVG formats. Final SVG and PDF files were audited to ensure that the vector publication versions contained no embedded raster-image objects. Superseded analytical and figure-generation attempts were retained for provenance but were not used as the manuscript-facing evidence layer.

### Ethics statement

This study analyzed publicly available, de-identified population-genomic data from the 1000 Genomes Project and did not involve newly collected human participant samples, new biological specimens, intervention, or direct participant contact. The source genomic data used for this secondary analysis were accessed for research purposes on 28 May 2026. The authors accessed only publicly available, de-identified genomic data and did not have access to names, contact details, or other information that could identify individual participants during or after data acquisition. No new institutional ethics approval was obtained for this secondary computational analysis because no new participants were recruited and no identifiable private information was accessed. The study cites and respects the original source-data consent, governance, and access framework associated with the 1000 Genomes Project.

### Use of artificial intelligence tools

OpenAI ChatGPT was used to assist with development and verification of analysis scripts, manuscript organization, language editing, and submission-formatting checks. AI-assisted outputs were checked against the author-reviewed analytical results, source-data documentation, reproducibility records, and cited literature. No primary data were generated or altered by the tool. Reuben S. Maghembe reviewed, tested, and revised all affected code and text and takes responsibility for the analyses, interpretations, and AI-assisted portions of the manuscript. All three authors reviewed and approved the final manuscript.

### Data and code availability

The source human genomic data are publicly available through the 1000 Genomes Project (1000 Genomes Project Consortium 2015). The analysis used the 2019-03-12 integrated, phased, biallelic SNV and INDEL GRCh38 call set described by Lowy-Gallego et al. (2019); target and control genotypes were derived from this same source release for the retained 660-person population panel.

The current Restrictome-EVOLVE analytical scripts, derived reproducibility materials, supplementary tables, figure-source data, and final publication assets for the population-resolved phased-haplotype and matched-genomic-control analysis are publicly available from the project repository at https://github.com/rmaghembe1/restrictome-evolve, with the manuscript-associated v2.0.1 release at https://github.com/rmaghembe1/restrictome-evolve/releases/tag/v2.0.1, and are archived in Zenodo at https://doi.org/10.5281/zenodo.21908890. Large primary 1000 Genomes VCF files are not redistributed; the source genomic data remain available through the 1000 Genomes Project as described above.

## Acknowledgments

The authors acknowledge the 1000 Genomes Project for making population-genomic data available to the research community. The authors also acknowledge the open-source scientific-computing ecosystem used to support reproducible genomic analysis, validation, visualization, and manuscript preparation.

## Abbreviations

ACB: African Caribbeans in Barbados
ASW: Americans of African Ancestry in SW USA
ESN: Esan in Nigeria
FDR: false discovery rate
FST: fixation index
GWD: Gambian in Western Divisions in The Gambia
LWK: Luhya in Webuye, Kenya
MAD: median absolute deviation
MSL: Mende in Sierra Leone
YRI: Yoruba in Ibadan, Nigeria.

## Supporting information captions

S1 Table. Canonical restriction-factor units and gene assignments.

S2 Table. Complete global endpoint/context comparative-inference results.

S3 Table. Global FDR-significant endpoint/context signals.

S4 Table. Pairwise population-differentiation results.

S5 Table. Population-private haplotype-structure results.

S6 Table. Within-population haplotype diversity and concentration results.

